# Sleep deprivation rapidly upregulates serotonin 2A receptor expression via the immediate early gene *Egr3*

**DOI:** 10.1101/634410

**Authors:** X. Zhao, K. T. Meyers, A. McBride, K. K. Marballi, A. M. Maple, K. L. Beck, P. Kang, M. Palner, A. Overgaard, G. M. Knudsen, A. L. Gallitano

**Affiliations:** Department of Basic Medical Sciences, University of Arizona College of Medicine – Phoenix, Phoenix, AZ, 85004; Interdisciplinary Graduate Program in Neuroscience, Arizona State University, Tempe, AZ, 85287; Arizona State University, Tempe, AZ, 85287; Epidemiology and Biostatistics, University of Arizona Mel and Enid Zuckerman College of Public Health – Phoenix, Phoenix, AZ, 85006; Department of Neurology and Neurobiology Research Unit Copenhagen University Hospital, Copenhagen, Denmark

## Abstract

Serotonin 2A receptors (5-HT_2A_Rs) mediate the effects of hallucinogenic drugs and antipsychotic medications, and are reduced in schizophrenia patients’ brains. However, the mechanisms that regulate 5-HT_2A_R expression remain poorly understood. We show that an environmental stimulus, sleep deprivation, upregulates 5-HT_2A_Rs in the mouse frontal cortex (FC) in just 6-8 hours. This induction requires the immediate early gene transcription factor early growth response 3 (*Egr3*). Further, EGR3 binds to the *Htr2a* promoter in the FC *in vivo*, and drives reporter construct expression *in vitro* via two *Htr2a* promoter binding sites. These findings suggest that EGR3 directly regulates FC *Htr2a* expression in response to physiologic stimuli, providing a mechanism by which environment rapidly alters levels of a brain receptor that mediates symptoms, and treatment, of mental illness.

**One Sentence Summary:** Just 6-8 hours of sleep deprivation upregulates brain levels of the receptor that mediates the response to hallucinogens.

---

Serotonin 2A receptors (5-HT_2A_Rs) mediate hallucinogenic effects of drugs including psilocybin, mescaline, and LSD (reviewed in (*1, 2*)), and are a key target of “second-generation antipsychotic” medications (SGAs) used to treat schizophrenia and other psychotic disorders (*3, 4*). Numerous studies have revealed that 5-HT_2A_R levels are reduced in the brains of schizophrenia patients, both post-mortem and *in vivo* ((*5–9*) and citations in (*10*)). However, the molecular mechanisms that regulate expression of this receptor that plays a critical role in the symptoms, and treatment, of mental illnesses remain poorly understood.

The delayed effect of medications that treat psychiatric illness have led to the concluion that the neurobiological changes that underlie their therapeutic response, occur in a timeframe of weeks. However, our results demonstrate that the environmental stimulus of sleep-deprivation, which produces rapid antidepressant effects in humans, upregulates 5-HT_2A_R levels in the mouse frontal cortex (FC) within just 6-8 hours (for mRNA and protein, respectively).

Our prior studies revealed that mice lacking the immediate early gene (IEG) *Egr3* have reduced levels of 5-HT_2A_R and, like humans experiencing psychosis, are resistant to sedation by SGAs, a feature shared by 5-HT_2A_R knockout mice (*11–14*). These findings led us to hypothesize that EGR3, an activity-dependent transcription factor, may directly regulate expression of the 5-HT_2A_R gene (*Htr2a*). If this were the case, it would suggest that *Htr2a*, like *Egr3*, should be upregulated in response to environmental stimuli.

Indeed, we found that 6h of sleep deprivation (SD), which upregulates *Egr3* in the cerebral cortex (*15*), significantly increased *Htr2a* mRNA in mouse cerebral cortex, and that this induction required *Egr3* (*16*). However, it was unclear whether this upregulated expression resulted in an increase in 5-HT_2A_R protein and, if so, whether this was occurring throughout the cortex, or in regions consistent with antero-posterior gradient in which *Htr2a* is expressed (*17, 18*).

We first tested whether we could replicate published *in situ* hybridization findings showing that SD upregulates *Egr3* (*15*), using quantitative reverse transcription (qRT) PCR. Figure 1A shows the SD protocol and coordinates for regional brain dissection. In WT mice, we found that 6h of SD did not increase *Egr3* expression in the most anterior region of frontal cortex (AFC), but did significantly upregulate *Egr3* mRNA in the posterior part of the frontal cortex (PFC), as well as in more posterior regions of cortex (labeled “mid-posterior cortex (MPC)) (Fig.s 1B – 1D).

**Fig. 1.**
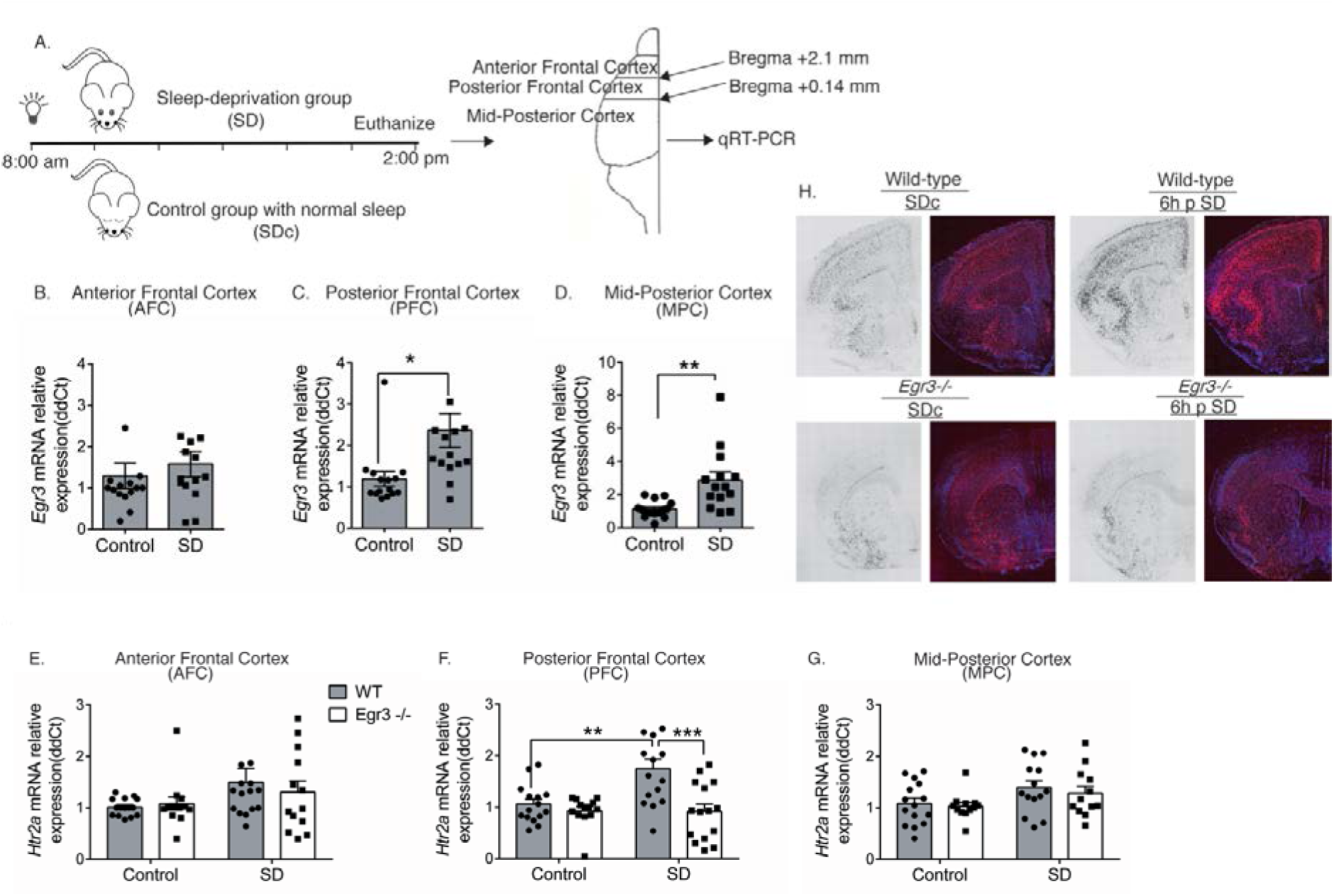
Sleep deprivation upregulates *Htr2a* in an *Egr3*-dependent, and region-specific, manner. (**A**) SD protocol. In WT mice quantitative RT-PCR shows that 6h of SD (**B**) does not increase *Egr3* expression in AFC regions (t_27_ = 0.6679; *p* = 0.510), but significantly upregulates *Egr3* mRNA in (**C**) PFC (t_28_ = 2.615, *p* = 0.0142) and (**D**) MPC (t_27_ = 3.5; *p* = 0.0016) regions, compared to SDc. (**E** – **G**) In WT and *Egr3*-/- mice 6h of SD (**E**) does not increase *Htr2a* expression in AFC (ANOVA; no sig. main effect of SD (F_1, 52_ = 3.488, p = 0.0675) or genotype (F_1, 52_ = 0.1129, p = 0.7382)), yet (**F**) significantly upregulates *Htr2a* expression in the PFC of WT, but not *Egr3*-/-, mice, compared to SDc (ANOVA: sig. main effect of SD (F_1, 54_ = 5.857, *p* = 0.0189), and sig. main effect of genotype (F_1, 54_ = 11.95, *p* = 0.0011); post-hoc analyses showed a sig. increase of *Htr2a* mRNA after SD (vs. SDc) in WT mice (*p* <0.01), but not in the *Egr3* -/- mice (*p* = 0.9998)). (**G**) In the MPC, SD increased *Htr2a* overall when both genotypes were analyzed (ANOVA, sig. main effect of SD (F_1, 49_ = 5.976, *p* = 0.0181)) but *Htr2a* mRNA increases were not sig. when comparing either genotype alone (post-hoc analyses showed no sig. differences between genotypes or SD conditions). (**H**) RNAscope *in situ* hybridization demonstrating *Htr2a* expression in SDc and SD WT and *Egr3*-/- mice (sections from PFC region). Bonferroni-corrected comparisons: * *p* < 0.05, ** *p* < 0.01, *** *p* < 0.001, *n* = 12-15. Values represent means ± SEM. (AFC: anterior frontal cortex; h: hours; PFC: posterior frontal cortex; SD: sleep deprivation; SDc: SD control; sig: significant, MPC: mid to posterior cortex; WT: wildtype).

We next examined whether the same 6h of SD can upregulate *Htr2a* expression in the same cortical regions, and whether this requires *Egr3* (Fig.s 1E – 1G). In the AFC, SD did not significantly increase *Htr2a* expression in either WT or *Egr3-/-* mice (Fig. 1E). However, in the PFC of WT mice, SD significantly increased *Htr2a* mRNA compared to SDc, a result not seen in *Egr3*-/- mice (Fig. 1F). In the MPC, SD increased *Htr2a* mRNA when both WT and Egr3-/- mice were analyzed together, but the increase was not significant in either genotype alone (Fig. 1G). RNAscope *in situ* hybridization shows that *Htr2a* expression is significantly lower in *Egr3*-/- than WT mouse PFC at baseline (SDc), and is not increased by 6h SD in *Egr3*-/- mice, as it is in WTs (Fig. 1H).

To determine if SD also increases 5-HT_2A_R protein expression, we performed receptor autoradiography in brain sections with 3H-M100907, a selective 5-HT_2A_R antagonist, following 8h SD (to allow time for translation of mRNA) (Fig. 2A). In the AFC, SD does not significantly increase 5-HT_2A_R binding in either WT or *Egr3*-/- mice (Fig. 2B). However, SD increases WT expression sufficiently to produce a significant difference in 5-HT_2A_R levels between WT and *Egr3*-/- mice that is not present in SDc animals.

**Fig. 2.**
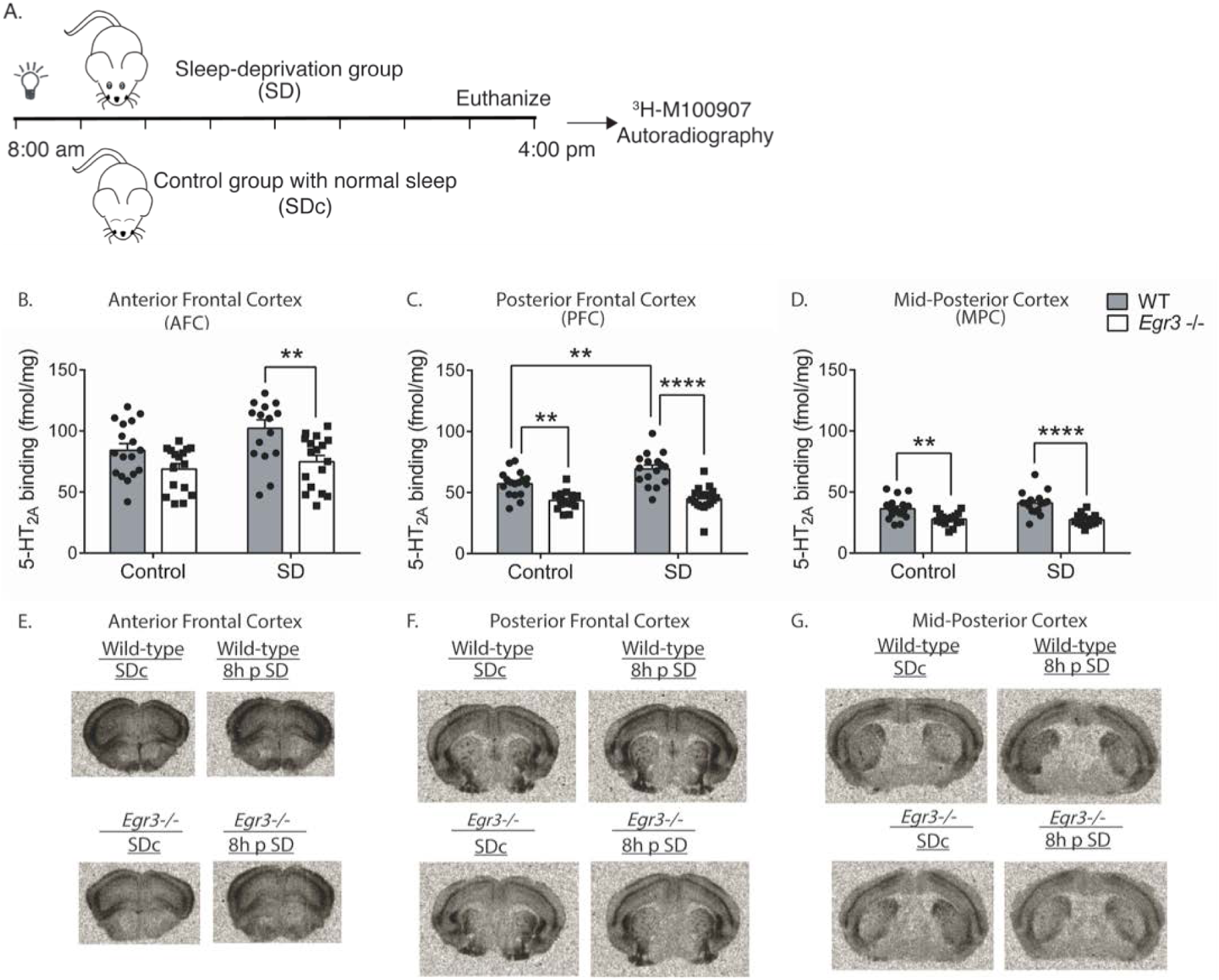
SD increases 5-HT_2A_R levels in the PFC of WT mice in an *Egr3*-dependent manner. (**A**) 8h SD protocol. Quantification of ^3^H-M100907 binding autoradiography shows that SD, compared with SDc: (**B**) in AFC results in significantly greater 5-HT_2A_R levels in WT mice than *Egr3*-/- mice after SD (ANOVA: sig. main effects of SD (F(1,62) = 4.61, p = 0.036) and genotype (F(1,62) = 14.78, p = 0.0003); (**C**) in the PFC SD significantly upregulates 5-HT_2A_R levels in WT, but not *Egr3*-/-, mice (ANOVA: sig. interaction between SD and genotype (F(1,62) = 4.18, p = 0.045). (**D**) In the MPC, SD did not significantly increase 5-HT_2A_R levels; notably, 5-HT_2A_Rs were lower in *Egr3*-/- mice than WT under both basal (SDc) and SD conditions (ANOVA: sig. main effect of genotype (F(1,62) = 38.79, p < 0.0001) but not of SD (F(1,62) = 1.371, p = 0.25)). Representative ^3^H-M100907 autoradiography images of brain tissue sections from (**E**) AFC, (**F**) PFC, and (**G**) MPC. Bonferroni-corrected comparisons: * *p* < 0.05, ** *p* < 0.01, *** *p* < 0.001, *n* = 16-17. Values represent means ± SEM.

In the PFC, 8h of SD significantly increases 5-HT_2A_R levels in WT mice but not in *Egr3*-/- mice (Fig. 2C). In addition, 5-HT_2A_R levels are significantly greater in WT than *Egr3*-/- mice in this region both at baseline (replicating our prior radioligand binding assay findings (*12*)), and following SD.

In the MPC, where endogenous *Htr2a* expression is lower than in more anterior cortical regions, SD does not increase 5-HT_2A_R levels in WT or *Egr3*-/- mice (Fig. 2D). Figures 2E – 2G show representative autoradiographic images from WT and *Egr3*-/- mice under SDc and SD conditions.

In all regions, radioligand binding reveals significant differences in 5-HT_2A_R levels between WT and *Egr3*-/- mice after SD and, in all but the most anterior (AFC) region, also at baseline (SDc). In contrast, differences in *Htr2a* mRNA levels between WT and *Egr3*-/- mice are seen only in the PFC region following SD. This difference could be due to the longer perdurance of protein than mRNA.

These data reveal the novel finding that 5-HT_2A_Rs can be upregulated in the PFC in a matter of hours in response to an environmental stimulus, and that this requires *Egr3*. These results suggest that EGR3, an activity-dependent IEG transcription factor, may directly regulate expression of *Htr2a* in response to environmental events.

We have previously shown that EGR3 is expressed in *Htr2a*-expressing neurons in the mouse FC, an essential criterion for EGR3 to potentially directly regulate *Htr2a* expression (*16*). Bioinformatic analyses identified two high probability EGR binding sites in the Htr2a promoter (Fig. 3A).

**Fig. 3.**
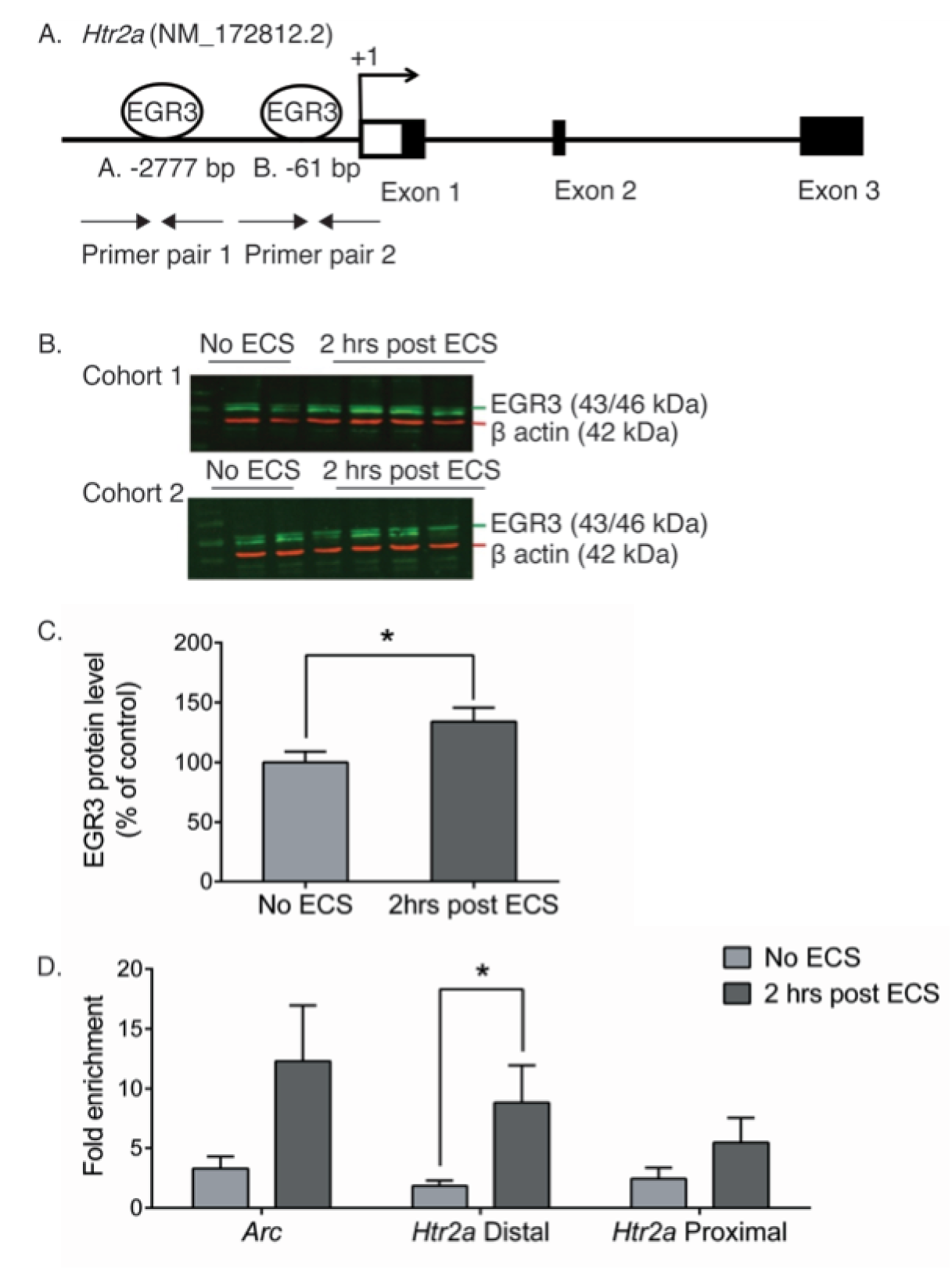
EGR3 binds to the *Htr2a* promoter in frontal cortex. (**A**) Schematic showing high probability EGR3 consensus binding sites in the *Htr2a* promoter. Western Blot **(B)** images and (**C**) average protein levels, show significant upregulation of activity dependent EGR3 protein 2h after electroconvulsive stimulation (ECS) in WT frontal cortex (n = 6). **(D)** ChIP-qPCR shows ECS increases binding of EGR3 to *Htr2a* distal promoter in frontal cortex issue (n = 11). Unpaired student *t*-tests, * *p* <0.05. Values represent means ± SEM.

To determine whether EGR3 protein binds to these binding sites in mouse cortex we conducted chromatin immunoprecipitation (ChIP). We used electroconvulsive seizure (ECS) to induce neuronal activity and maximally express EGR3 in the PFC (Fig.s 3B and 3C). Compared to the positive control region, the promoter of activity-regulated cytoskeleton associated protein (*Arc*) (a validated EGR3 target gene (*19*)), binding of EGR3 to the distal *Htr2a* promoter was significantly increased following ECS, compared to non-stimulated controls (Fig. 3D).

To confirm that the binding of EGR3 to the *Htr2a* promoter results in a change in gene expression, we conducted *in vitro* luciferase-reporter assays. We co-transfected neuro2a cells with luciferase/SEAP constructs driven by either the positive control *Arc* promoter (*19*), the distal *Htr2a* promoter, or the proximal *Htr2a* promoter, with either a CMV vector overexpressing EGR3, or a control empty CMV vector (Fig. 4A-C). We found that both regions of the *Htr2a* promoter containing high-probability EGR3 binding sites (Fig. 3A) drive expression of luciferase in response to EGR3 expression. EGR3 expression induces a 4.9 fold increase in the positive control *Arc* promoter-driven luciferase (Fig. 4A) and 3.9-fold increase in the *Htr2a* distal promoter-driven luciferase (Fig. 4B). In addition, although the proximal *Htr2a* promoter did not show a statistically significant increase in EGR3 binding in the ChIP assay, *in vitro* expression of EGR3 induced a 4.2-fold increase in *Htr2a* proximal promoter-driven luciferase signal, compared to CMV vector alone (Fig. 4C). These results suggest that EGR3 directly binds to the *Htr2a* promoter in the cortex in response to neuronal activity, and activates *Htr2a* expression, which results in increased levels of cortical 5-HT_2A_Rs.

**Fig. 4.**
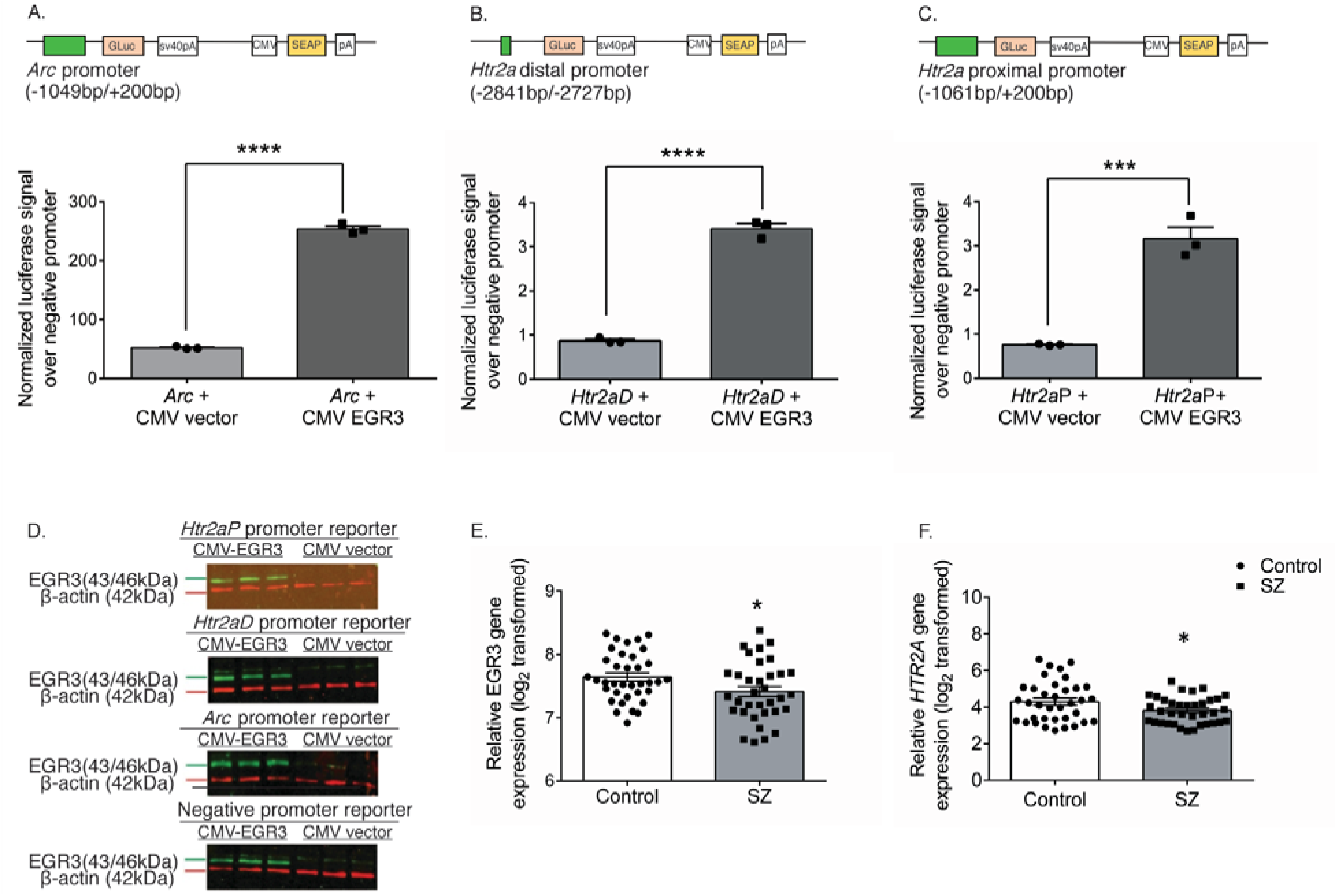
EGR3 drives gene expression via binding sites in the *Htr2a* promoter. (**A-C**) Schematics of dual luciferase/SEAP reporter constructs containing EGR consensus binding sites and results of *in vitro* assays in neuro2a cells. CMV-driven EGR3 overexpression significantly upregulates expression of luciferase reporters driven by (**A**) *Arc* promoter, (**B**) *Htr2aD* distal promoter (t4 = 21.17, p < 0.0001), and (**C**) *Htr2aP* proximal promoter (t4 = 8.977, p < 0.001) regions. (**D**) Western blot validation of EGR3 expression following transfection with CMV-EGR3 versus CVM empty vector, from cultures expressing reporter constructs driven by promoters from ARC, *Htr2aD*, *Htr2aP*, or negative promoter control vector. Unpaired student *t*-tests, \*\*\**p* < 0.001, \*\*\*\**p* < 0.0001, *n* = 3. (**E, F**) *EGR3* and *HTR2A* mRNA levels are significantly decreased in human brain tissue samples from the prefrontal cortex of schizophrenia patients compared to controls. Microarray (Robust Multi-Array Average) gene expression data derived from NCBI Geo database GSE53987 showing significant decrease in (**E**) *EGR3* (\**p* < 0.033) expression and (**F**) *HTR2A* (\*\**p* = 0.005) expression, in control (N = 19) vs. schizophrenia (N = 15) patients (Mann Whitney U test). Values represent means ± SEM. (Abbreviations: CMV - cytomegalovirus, GLuc - *Gaussia* luciferase, SEAP - secreted alkaline phosphatase.)

These results reveal a novel mechanism of 5-HT_2A_R regulation, rapid environmentally-induced expression of *Htr2a* via the activity dependent IEG transcription factor EGR3. Prior studies have shown that 5-HT_2A_R levels can be altered in response to ligand binding; for example, 5-HT_2A_R agonists and antagonists trigger receptor internalization and recycling (*20, 21*). However, little is known about transcriptional regulation of *Htr2a*, or that environmental stimuli rapidly alter 5-HT_2A_R levels. Interestingly, our findings are supported by a recent human study reporting that 24 h sleep deprivation causes a significant increase in brain 5-HT_2A_R levels detectably by Positron Emission Tomography (PET) scan (*22*).

5-HT_2A_Rs are abundantly expressed in the neocortex and play important roles in cognition and mood. They also mediate hallucinogenic effects of numerous drugs including LSD, psilocybin, and mescaline (*1, 2*). Investigation of these drugs has recently undergone a resurgence in the search for treatments for severe psychiatric symptoms including depression (*23*) and anxiety disorders including PTSD (*24–26*). 5-HT_2A_Rs are also a key target of SGAs, which treat the symptoms of psychosis (*4*).

As far back as 1976 numerous post-mortem and *in vivo* studies have revealed that 5-HT_2A_R (or 5-HT_2_R) levels are reduced in schizophrenia patients’ brains (*5–10*). Such findings may lead to the assumption that brain neurotransmitter receptor levels are a relatively stable characteristic.

Our findings suggest, in contrast, that levels of this critical receptor are dynamically, and rapidly, regulated at the transcriptional level in response to neuronal activity and in response to environmental events. This suggests the intriguing possibility that the reduced 5-HT_2A_R levels reported in schizophrenia patients may be a consequence of disrupted neural activity, an hypothesis supported by numerous findings of abnormal IEG expression in patients’ brains (Fig. 4E, (*27, 28*).

Although genome wide association studies (GWAS) have recently identified numerous loci believed to increase genetic risk for illnesses like schizophrenia, the mechanism by which environment may interact with these regions remains elusive. Notably, *EGR1, EGR4*, and *NAB2* (a transcriptional co-regulator that alters gene expression via binding to the EGRs) each map to one of the 145 GWAS loci for schizophrenia (*29*). Although *EGR3* itself is not within a GWAS locus, it interacts in co-regulatory feedback loops with the EGRs and NAB2 (*30, 31*), and *EGR3* expression is reduced in the brains of schizophrenia patients, including in our analyses of data from the NCBI GEO database, showing reduced levels of both *EGR3* and *HTR2A* mRNA in the prefrontal cortex of schizophrenia patients, compared with controls (Fig. 4E and 4F) (*27*).

These findings suggest that dysfunction in activity dependent EGR family IEGs, which include, and result in, decreased activity of EGR3, may contribute to the reported deficits in 5-HT_2A_R expression in schizophrenia patient brains. These findings thereby shed light on a potential mechanism whereby environment may interact with genetic variations to influence neurobiology that may contribute to the symptoms, and treatment, of neuropsychiatric illness.

## Acknowledgments

We are grateful to L Muppana, D Elizalde, and J Campbell for animal colony maintenance and technical assistance, to A. Aden, A. Barkatullah, M. Charbel, R. Khoshaba, E. Offenberg and A. Ozols for assistance with SD studies, to A. Bhaskara and M. Godbole for NCBI GEO data analyses, and to A. Gulledge for expert advice.

## Funding

This work was supported by NIH R01 award MH097803 (ALG).

## Author contributions

X. Zhao - Lead, Writing – original draft, Investigation, Formal analysis, Visualization; K. T. Meyers - Investigation, Formal analysis, Visualization; A. McBride – Investigation, Resources; K. K. Marballi - Investigation; A. M. Maple - Project administration; C. Raskin - Investigation; J. Lish - Investigation; A. Misra - Investigation; S. Noss - Investigation, K. L. Beck - Investigation; P. Kang - Formal analysis; M. Palner - Investigation; A. Overgaard - Investigation; G. M. Knudsen - Supervision; A. L. Gallitano - Conceptualization, Funding acquisition, Supervision, Writing – review & editing.

## Competing interests

Authors declare no competing interests.

## Data and materials availability

All data is available in the main text or the supplementary materials. Data for Fig. 4E-F are available from NCBI GEO database, Accession: GSE53987.

## Notes

#### Summary of Updates

Addition of authors, edits to figures, condensation and revision of text.

