## Supplemental methods and figure for "Sleep deprivation rapidly upregulates serotonin 2A receptor expression via the immediate early gene *Egr3*"

**Materials and Methods**

Animals

*Egr3*-/- mice were generated by the deletion of sequences that encode the zinc fingers, the DNA-binding domain of the protein *(32)*. The mice were backcrossed to a C57BL/6 background for more than 30 generations. Animals were maintained as heterozygote × heterozygote mating pairs. *Egr3*-/- and their WT littermates were assigned as 'matched pairs' at the time of weaning. Male adult littermates were used and housed on a 14/10 h light/ dark schedule with ad libitum access to food and water.

Sleep deprivation (SD)

Animals were single-housed for 5 days prior to the experimental procedure for habituation. For gene expression level study, 6hrs of SD was performed in two groups of animals (Fig.S1): 6 hrs SD group, time - matched controls allowed to sleep in their home cages (control group). For radioligand binding of protein level study, 8hrs of SD was performed in two groups of animals (Fig.S1): 8 hrs SD group, time - matched controls allowed to sleep in their home cages (control group).

SD started at the beginning of the light period (8:00 a.m. mountain standard time, MST). Mice were kept awake by "gentle handling" as previously described *(15)*. Briefly, animals were disturbed by a combination of cage tapping, introduction of novel objects (e.g., balled paper towels), cage rotation, and stroking of vibrissae and fur with an artist's paintbrush *(15)*. The number and type of stimulation required to keep each animal awake during the SD procedure were recorded and compared between the WT and *Egr3*-/- mice.

Quantitative reverse transcription polymerase chain reaction (qRT-PCR)

Animals were sacrificed immediately after SD via isoflurane overdose. The brains were removed and the following regions were immediately dissected: the anterior frontal cortex, posterior frontal cortex, and the mid-posterior cortex. Collected cortical tissues were treated with RNAlater solutions (ThermoFisher Scientific, Waltham, MA) for ribonucleic acid (RNA) stabilization and storage. For RNA isolation, the tissue was homogenized in 700μl of TRI reagent (Life Technologies, Carlsbad, CA) in the 2ml tubes (Bertin Corp, Rockville, MD) prefilled with ceramic beads (diameter 1.4 mm). Tissue was homogenized for 4 cycles at a speed of 6000 g for 30 s, in a Precellys 24 high-powered bead mill homogenizer (Bertin Corp, Rockville, MD). The samples were put on ice for 2 minutes between each cycle to prevent RNA degradation. Next, the homogenates were centrifuged at 12000g, 4°C for 5 min. The supernatant was saved and was separated into aqueous and organic phases by adding 70μl per sample bromochloropropane (BCP, Molecular Research Center, Inc. Cincinnati, OH). The RNA was precipitated in a MagMAX express magnetic particle processor (Applied Biosystems, Foster City, CA) with isopropanol and washed with ethanol. Next, RNA was dissolved in nuclease-free water and quantified with ND-1000 Spectrophotometer (Thermo Scientific, Waltham, MA) and further confirmed with Qubit 3.0 Fluorometer (Thermo Scientific, Waltham, MA). The mRNA was reverse transcribed into cDNA using M-MLV reverse transcriptase kit (Life Technologies, Carlsbad, CA). Quantitative RT-PCR was performed using FastStart SYBR green master mix (Roche Applied Science, Indianapolis, IN) with the reaction volume 20μl per well (10μl SYBR green master mix 2X, 8μl cDNA, 1μl of each forward and reverse primers of 10uM stock) in a 7500 Fast real-time PCR machine (Applied Biosystem, Foster City, CA). The thermal cycling parameters are 50 °C for 2 minutes, 95 °C for 10 minutes, 40 cycles of (95 °C for 15 seconds, 60

°C for 1 minutes). Quantitative RT-PCR primers were designed using primer3 *(33)* and target specificity was confirmed using the software primer-BLAST (https://www.ncbi.nlm.nih.gov/tools/primer-blast/). Primer sequences were ordered from Integrated DNA Technologies (IDT, Coralville, IA) with standard desalting purification. Primers used for the study are listed as below:

*Egr3* Forward primer: 5'-CTGGAGGCACTTTCTCCTTG-3'
*Egr3* Reverse primer: 5'-TGGATCAAGGCGATCCTAAC-3'
*Htr2a* Forward primer: 5'-TCACCATTGCGGGAAACAT-3'
*Htr2a* Reverse primer: 5'-ATCAGCTATGGCAAGTGACAT-3'
*Pgk1* Forward primer: 5'-TGTTAGCGCAAGATTCAGCTA-3'
*Pgk1* Reverse primer: 5'-CAGACAAATCCTGATGCAGTA-3'

<u>*In situ* hybridization</u>

Age matched, paired WT and *Egr3*-/- female mice (3-5 mo. old) underwent 6 hours of sleep deprivation. Mice were sacrificed and tissue was flash frozen. 12 micron thick tissue sections from WT and *Egr3*-/- pairs that underwent sleep deprivation were mounted on the same slide as their non-sleep deprived controls. Tissue was stored in the -80C until the day of the experiment.

Tissue was removed from the freezer on the day of the experiment and immediately transferred to freshly prepared 4% paraformaldehyde solution at 4°C for 1 h. Tissue was taken through a gradient of alcohol dehydrations and was dried before applying the hydrophobic barrier around the tissue sections.

*In situ* hybridization was performed using RNAscope technology (ACD a Biotechne Brand), following the RNAscope Multiplex Fluorescent Reagent Kit v2 User Manual (Document Number 323100-USM) with the RNAscope Multiplex Fluorescent Reagent Kit V2 (323100). A probe designed against *Htr2a* was purchased from Advanced Cell Diagnostics, Inc (Probe-Mm-Htr2a-C3, 401291-C3) and was developed with the fluorescent Opal dye 570 (FP1488001KT, PerkinElmer) at a concentration of 1/750. Tissue was counterstained with DAPI (Vectashield Antifade Mounting Media with DAPI, H-1200).

<u>Tissue Collection/Sectioning for radioligand binding study</u>

Male WT and *Egr3*-/- mice (aged 55-62 days) were anesthetized with isoflurane and sacrificed. Tissue was flash frozen in -40°C methylbutane and stored at -80°C. Tissue was sectioned at a thickness of 12 microns on a cryostat. Six sections of anterior frontal cortex (Bregma 2.8 – Bregma 2.34), posterior frontal cortex (Bregma 1.70 – Bregma 0.86), two sections of mid-cortex (Bregma 0.02 - -0.82), and two sections of posterior cortex (Bregma -1.34 - -2.30) were collected and freeze-thawed on the same slide. A total of ten slides/brain were serially collected and stored at -80°C.

<u>Radioligand Binding Analysis - Autoradiography</u>

Tissue was treated with 3-H-MDL100907 [R(+)-α-(2,3-dimethoxyphenyl)-1-[2-(4-fluorphenyl)-ethyl]-4-piperidin-methanol] (specific activity; 59.22 Ci/mmol), a selective 5HT2A antagonist, and was detected with autoradiography. Control tissue lacking drug was exposed to assess non-specific binding.

Autoradiography Analysis

Standards of known activity were included in each cassette. Treated and non-treated tissue was exposed within the same cassette with the standards. Mean intensity values of the standards were determined by outlining the standard curve regions with the Quantity One, Bio-Rad program (Bio-Rad Laboratories, Hercules, CA). A derived equation of the standard curve was generated from standard mean intensity values. The mean intensity values of ligand-bound cortical and hippocampal regions of interest were generated by free hand contour tracing. Ligand-bound (fmol/mg) measurements were calculated from mean intensity values of the regions of interest extrapolated from the derived equation, divided by known activity of the drug. Mean intensity values for tissue with non-specific binding was determined and ligand bound values were extrapolated. Specific ligand bound activity was subtracted from non-specific activity to obtain corrected values.

Image processing

RNAscope images were taken at a 20x magnification with Zeiss Axio Imager M2 epifluorescent microscope in each individual channel (DAPI, 3.2 ms and Cy3, 300 ms). Post-processing of images was conducted in an identical fashion in Adobe Photoshop by adjusting the images taken in channel Cy3 to a brightness of 100, contrast of 50, and white/black balance levels to 135 and 255, the exposure to 2.65 and the offset to -0.0059. Images taken in the DAPI channel were processed by adjusting white/black balance to 0 and 50, respectively. Images taken from the same section/animal of the DAPI/Cy3 were merged with ImageJ Fiji software. A separate post-processing was conducted solely for the images taken in the Cy3 channel in Adobe Photoshop by inverting the image and adjusting the white/black balance levels to 135 and 255, respectively. Images were then saved in the Red channel of Adobe Photoshop.

Autoradiography images were acquired from the original exposed Cassette image (.tiff) by opening the image in Adobe Photoshop, and cropping the images of the individual tissue sections at the same magnification (200x). Images were saved and imported into Adobe Illustrator. Each image underwent adjustment of white/black balance levels 65 and 200, respectively, were inverted, and adjusted with a brightness of 125 and a contrast of -50.

Electroconvulsive seizure (ECS)

Ten minutes after placing a drop of proparacaine hydrochloride ophthalmic solution USP, 0.5% (Akorn) onto each eye, the electroconvµlsive shock was delivered via silver trans - corneal electrodes previously damped with 0.9% sodium chloride (NaCl, Sigma-Aldrich). Animals were restrained by manual scruffing. The pulse generator (ECT Unit 57800 - 001; Ugo Basile, Comerio, Italy) delivered a stimulus of 220 - 250HZ (based on weight), 0.9 width square wave pulses for a duration of 0.2 seconds at a current of 20mAs. Immediately following the shock, mice displayed tonic-clonic seizures with hind limb tonic extensions and were placed back in their home cages for recovery. Mice were sacrificed and their brains were dissected 2 hrs after ECS. Age-matched “no-ECS” control mice from the same litters were sacrificed at the same time of day as the mice experiencing seizures.

Western blot to detect the expression pattern of EGR3 protein 2hrs after ECS

2 hours after ECS, posterior frontal cortex was removed and homogenized in the 1% Nonidet P-40 lysis buffer containing 0.5% sodium deoxycholate with proteinase inhibitor (1:10,000; Sigma-Aldrich, St. Louis, MO) using the Wheaton Tenbroeck style tissue grinder

(ThermoFisher Scientific, Waltham, MA). Cell debris was pelleted by centrifugation at 17,000 rpm for 5 min, 4 °C. The supernatant was saved and protein concentration was quantified using Pierce BCA protein assay kit (ThermoFisher Scientific, Waltham, MA). Protein samples were denatured for 5 min, 100 °C on block heater (VWR, Radnor, PA). 25ug of protein was loaded into homemade 4-8% gradient polyacrylamide gels. Following electrophoresis, proteins were transferred to nitrocellulose membranes (Bio-Rad Laboratories, Hercules, CA) for 1hr using a semi-dry transfer apparatus (Bio-Rad Laboratories, Hercules, CA) which constantly delivers a voltage of 15 V. Then the membranes were blocked for 1hr with 3% (w/v) non-fat dry milk (LabScientific Inc., Highlands, NJ). After washing with phosphate-buffered saline plus 0.1% of Tween-20 (PBST), the membranes were incubated overnight at 4°C with rabbit anti-EGR3 antibody (1:200, Santa Cruz Biotechnology, Dallas, TX) and mouse anti-beta actin antibody (1:5000, Sigma-Aldrich, St. Louis, MO). The next day, membranes were washed with PBST and incubated with IRDye800CW secondary antibodies (1:10,000 for EGR3, Li-COR Biosciences, Lincoln, NE) and the IRDye680 secondary antibodies (1:20,000 for beta actin, Li-COR Biosciences) dissolved in 3% non-fat dry milk for 1hr at room temperature with gentle shaking. After washing for 5 times, each time for 5 minutes, the membranes were visualized on an Odyssey instrument (Li-COR Biosciences). Protein expression levels were determined as the ratio of EGR3 to the internal control beta-actin and were reported as percentage of the control group.

Bioinformatics analysis to identify putative EGR3 binding sites on the *Htr2a* gene promoter

To identify potential EGR3 binding sites on the *Htr2a* gene promoter we downloaded the promoter sequences, including the region 4kb upstream of the *Htr2a* transcription start site (NM_172812, chr14:74636840-74640839) from genome browser of the University of California, Santa Cruz (UCSC genome browser, https://genome.ucsc.edu/). Then we scanned and identified matches of consensus binding sites for the EGR3 in the 4kb *Htr2a* promoter sequences using the software ‘Find Individual Motif Occurrences’ (FIMO, http://meme-uite.org/tools/fimo) *(34)*. The motif occurrences with a p value less than 0.0001 were selected.

Chromatin immunoprecipitation (ChIP)

2hrs after ECS, frontal cortex was removed and cut into 1mm pieces. The protein-DNA complexes of the frontal cortical tissue were crosslinked through incubation with 1% formaldehyde (Sigma-Aldrich, St. Louis, MO) on a rotator for 12 minutes. Formaldehyde was quenched with 125 mM glycine (Sigma-Aldrich, St. Louis, MO) for 5 minutes. Samples were homogenized using a Q125 sonicator (Qsonic, LLC, Newtown, CT) with a power of amplitude 40%, sonication duration 7, for 2 cycles. Samples were cooled on ice between each cycle. Chromatin was then sheared into 200bp-1000bp with a Bioruptor XL (Diagenode Inc., Denville, NJ) at 4°C at a sonication intensity for 30s on/30s off for 35 cycles. Fragment size was verified with an Agilent bioanalyzer (Agilent Technologies, Santa Clara, CA). 50μl of sheared chromatin were removed as input control with an input fraction of 5%. The magnetic beads-antibody complex was prepared by incubating 7.5ug of the anti-EGR3 antibody (Santa Cruz Biotechnology, Dallas, TX) with the magnetic sheep anti-rabbit beads (Invitrogen Corp., Carlsbad, CA) at 4°C overnight on a rotator. After washing the bead-antibody complex with 0.5% BSA blocking solution, 70μl of the beads-antibody complex was added into each ChIP sample, and incubated for 16hrs at 4°C on a rotator. To control for nonspecific binding, a normal IgG IP was performed in parallel. Beads were collected in sample tubes with a magnetic rack

(ThermoFisher Scientific, Waltham, MA), and washed with the following buffer: low salt wash buffer (0.1%SDS, 1% TritonX100, 2mM EDTA, 150mM NaCl, 20mM Tris-HCl), high salt wash buffer (adjusted with 200 mM NaCl in place of the concentration of NaCl of the low salt wash buffer), LiCl wash buffer (150mM LiCl, 1%NP40, 1% NaDOC, 1mM EDTA, 10mM Tris-HCl). Cross-linked protein and DNA complex were reversed at 65°C overnight. RNA was stripped by 1hr incubation with 2µl of RNase A (Roche Applied Science, Indianapolis, IN) at 37°C. Proteins were digested with 2µl of proteinase K (20 mg/mL, Invitrogen Corp., Carlsbad, CA). DNA was purified with a DNA purification kit (QIAGEN Inc., Germantown, MD). qPCR was performed using FastStart SYBR green master mix (Roche applied science) in a 7500 Fast real-time PCR machine (Applied Biosystem, Foster City, CA). The total reaction volume per well was 25µl reaction (2µl DNA, 9.5 µL nuclease-free water, 12.5 µL SYBR-Green Master Mix 2X, 0.5µl µL of each forward and reverse primers with 10uM stock). Thermal cycling parameters were designated as initial denaturation at 94°C for 10 minutes, 50 cycles of (denature at 94°C for 20 seconds, anneal and extension at 60°C for 1 minutes). The primers used are listed as follows:

*Arc* forward: 5'-TCGCTGCCCAGGACTAGGTA-3';
*Arc* reverse: 5'-TTCACAGCCCCGAGTGACTAA-3';
*Htr2a* proximal forward: 5'-CTTGGATAGAAGTGCTGGATGCT-3';
*Htr2a* proximal reverse: 5'-GGGTACATGGCAGTCATATTTTTAGG-3';
*Htr2a* distal forward: 5'-CTGGGCTCTAAAGGCAACTGA-3';
*Htr2a* distal reverse: 5'-TGCGCACGTGTATACAGAGTAGGT-3'

After performing ChIP- qPCR, the relative occupancy (aka. fold enrichment) of the EGR3 proteins at predicted binding loci of *Htr2a* putative regulation regions is estimated using the following equation 2^(ΔCt MOCK- ΔCt SPECIFIC), where ΔCT MOCK and ΔCT SPECIFIC are mean normalized threshold cycles of PCR done in triplicate on DNA samples from MOCK (anti-IgG antibody) and transcription factor EGR3 immunoprecipitations to the input IPs.

<u>Promoter reporter vector design</u>

The *Htr2a* proximal promoter luciferase reporter – the dual-reporter system contains a Gaussia luciferase gene (GLuc) which is driven by a *Htr2a* proximal promoter insert which corresponds to *Htr2a* promoter sequence located approximately 1061bp upstream and 200 bp downstream of the transcription start site (TSS) of the *Htr2a* gene. In the same vector, a secreted alkaline phosphatase (SEAP) is driven by a cytomegalovirus (CMV) promoter and serves as the internal control for signal normalization. This internal control SEAP exists in all the other promoter reporter clones in the present study. The *Htr2a* proximal promoter luciferase reporter contains the EGR3 putative binding site GCGCGGGGGAGGGG.

The *Htr2a* distal promoter luciferase reporter – the dual luciferase promoter reporter clone contains a GLuc gene and is driven by the insert, which is -2727 bp to -2841 bp upstream of the *Htr2a* TSS. This fragment contains the EGR3 putative binding site AGGAGGGGGAGTCT.

The *Arc* promoter luciferase reporter was used as a positive control for the illuminometer and the functionality of our CMV-EGR3 vector. We used the *Arc* promoter reporter clone containing an insert 1049bp upstream and 200bp downstream of the TSS of the *Arc* gene. This *Arc* promoter luciferase reporter contains the EGR3 binding site which was confirmed previously *(19)*.

The non-promoter luciferase reporter was used as a negative control to detect the basic activity of the dual-reporter vector. This luciferase reporter contains an insert that is a non-promoter sequence (TGCAGATATCCTCGCCC).

All the promoter clones were generated by Genecopeia (Genecopeia Inc., Rockville, MD).

Cell culture and transfection

Neuro2a cells (mouse neuroblastoma cells; ATCC, Manassas, VA) were maintained in Dulbecco's Modified Eagle Medium (DMEM, Gibco, Thermo Fisher Scientific) which contains 10% Fetal Bovine Serum (FBS, Gibco) and 1% Penicillin-Streptomycin (PS, Gibco), at 37°C, under a humidified atmosphere of 5% carbon dioxide (CO2) : 95% air. Cells were seeded into a 6-well plate (Corning Inc., Corning, NY) with 2 ml of cell culture medium and were transfected at the point of 70-90% confluence. Cell culture medium was removed 3 hrs before transfection and 3ml of fresh medium were added to each well. 1.5 ug of CMV-*Egr3* vector or CMV vector was co-transfected with 1 ug of each promoter reporter vector per well using Lipofectamine 3000 reagents (ThermoFisher Scientific). Transfection was performed with 3.75 µl of Lipofectamine 3000 reagent, 5 µl of P3000 reagent and 250 µl of Opti-MEM per well. Cells were incubated at 37°C with 5% CO2 : 95% air following transfection until further processing.

Three separate transfections were performed. Transfections were conducted in triplicate.

Luciferase signal measurement

24 hrs after transfection, 0.2 ml of medium from each cell culture was collected and placed at room temperature. The duo luciferase activities were measured using the secrete-pair dual luminescence assay kit (Genecopoeia). Each sample was run in duplicate.

To detect the Gaussia luciferase (GLuc) signal, the 10X Gaussia luciferase stable buffer (GLuc-S) was diluted with distilled water to 1X GLuc-S (1:10). The GLuc assay working solution was made by diluting the Gaussia luciferase substrate with the 1X GLuc-S buffer (1:10) and was incubated in the dark at room temperature for 25 minutes. 100 µl of GLuc assay working solution was mixed with 10 µl of cell culture medium for each well. The mixture was incubated at room temperature for 1 minute in the dark. The signal was read with a Tecan Safire2 instrument (Tecan Group Ltd., Morrisville, NC).

50 µl of each culture medium was aliquoted and heated at 65°C for 15 min, and placed on ice. 1X SEAP buffer was prepared from the 10X SEAP buffer working stock The SEAP assay working solution was diluted with the SEAP substrate with 1X SEAP assay working solution (1:10) and incubated at room temperature for 10 minutes in the dark. 100 µl of SEAP assay working solution was mixed with 10 µl of each heated medium sample. The mixture was incubated at room temperature for 10 minutes in the dark, Secreted alkaline phosphatase levels were then read with the Tecan Safire2 instrument (Tecan Group Ltd.).

For all measurements, the GLuc value was first normalized to the intern control SEAP luciferase value (GLuc / SEAP ratio) and then to the non-promoter luciferase reporter.

Western blot to measure the EGR3 protein level 24hrs after transfection

After collecting the cell culture medium for luciferase measurement, the remaining medium in each well was discarded. Cells were washed 2 times with cold 1X PBS, 300 µl per well and were lysed with 300 µl per well of 1% Nonidet P-40 lysis buffer containing proteinase inhibitor (1:10,000; Sigma-Aldrich). Samples were further homogenized using a Q125 sonicator (Qsonica) with a power of amplitude 40%, 1 time for 5 seconds on ice. Protein concentrations were quantified with a Nanodrop 1000 spectrophotometer (Thermo Scientific). Protein samples were denatured and 25ug of proteins for each sample were loaded on gels. The western blot protocol follows that of our previously described method for ECS.

NCBI GEO Data Analysis

We used publicly available data on NCBI GEO, Gene Expression Omnibus, a public repository for gene expression studies mainly using RNA sequencing and microarray data *(35)*. Data from dataset GSE53987 *(27)* were used in our study. We used the main microarray platform file GPL570 as a guide to ascertain gene -specific probe IDs for EGR3 and HTR2A. Subsequently, gene expression data for prefrontal cortex (Brodmann Area 46) EGR3 and HTR2A were downloaded from individual sample files by using the probe IDs as a reference to get the corresponding gene expression values. We collated the gene expression data for different EGR3 and HTR2A probes per sample per diagnostic group and did pairwise comparisons for each probe comparing schizophrenia vs. control samples. Of these, data for probes 211616_s_at (HTR2A) and 206115_at (EGR3) showed significant differences and are shown in the manuscript.

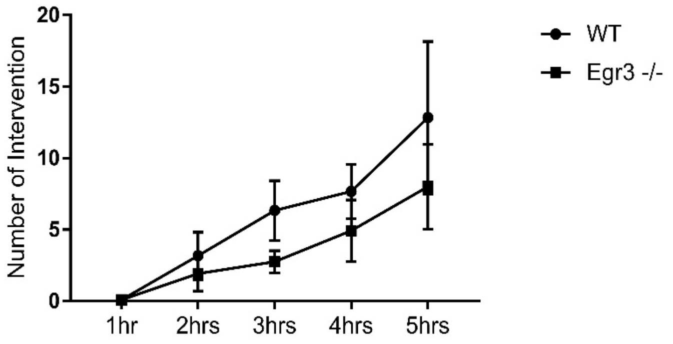


**Fig. S1.**

**WT and Egr3 -/- mice require same amount of stimulus during SD.** No significant effect of interaction between time and genotype is observed ($p = 0.8411$). A significant time effect ($p < 0.001$) but no significant genotype effect on the amount of stimulus ($p = 0.1461$) is detected. Two - way ANOVA with repeat - measurement (RM). n = 12. All values included in the figure legends represent means ± SEM.
